# Automated registration of spatial expression data scales multimodal integration to large cohorts

**DOI:** 10.1101/2025.05.29.656851

**Authors:** Caitlin F. Harrigan, Ching Yeung Lam, Danian Chen, Christian Lai, Rod Bremner, Hartland W. Jackson, Kieran R. Campbell

**Affiliations:** Department of Computer Science, University of Toronto, Toronto, Ontario, Canada; Vector Institute, Toronto, Ontario, Canada; Lunenfeld-Tanenbaum Research Institute, Sinai Health System, Toronto, Ontario, Canada; Ontario Institute of Cancer Research, Toronto, Ontario, Canada; Department of Ophthalmology and Visual Science, University of Toronto, Toronto, Ontario, Canada; Department of Laboratory Medicine and Pathobiology, University of Toronto, Toronto, Ontario, Canada; Department of Molecular Genetics, University of Toronto, Toronto, Ontario, Canada; Department of Statistical Sciences, University of Toronto, Toronto, Ontario, Canada

**Keywords:** Spatial proteomics, Highly multiplex imaging, Cross-modality registration, Bayesian optimization

## Abstract

Recent advances in spatial proteomics enable quantification of the spatial distribution of protein expression across a variety of scales, resolutions, and multiplexing. Registering images from such technologies across modalities is an essential task that enables both the integration of complementary imaging technologies and validation of biological findings. This can be particularly challenging when the modalities capture fundamentally different types of data, such as light intensity, probe counts, or heavy metal counts. However, few datasets and methods address this problem at scale. Here, we introduce the largest dataset to date of cross-modality imaging of both cell line and tissue slides suitable for benchmarking registration methods. We further present Twocan, a Bayesian optimization framework that enables robust automated registration between immunofluorescence imaging and highly multiplex spatial proteomics data. Our method achieves significantly higher registration success rates compared to existing approaches across our comprehensive dataset of 954 image pairs.

## INTRODUCTION

Highly multiplexed spatial proteomics technologies ^1^, such as Imaging Mass Cytometry (IMC) ^2^, can quantify the expression of more than 40 proteins at subcellular resolution, enabling high-dimensional readouts of cells and their spatial context within tissues. In contrast, immunofluorescence (IF) technologies measure the expression of fewer proteins (1-3) at much higher resolution (0.01-0.*µm* compared to 0.5-10*µm*), enabling highly sensitive measurement of key biomarkers in tissue sections ^3^. The contrasting strengths and limitations of these imaging techniques means it is highly advantageous to integrate them together across modalities ^4–9^. Such integration is essential for validation of biological experiments, development of new analyses, and additionally useful for training machine learning models, both supervised and unsupervised ^10^.

However, integration of paired images relies on correct alignment to a global coordinate system. Several tools exist for manual registration ^11,12^ and as spatial proteomics datasets continue to grow to contain thousands of images ^3,13–15^, automated approaches to image registration are being pursued ^5,16,17^. However, most existing methods do not work directly on data from multiplexed imaging technologies, which typically spatially measure heavy metal or ion counts. As a result, substantial manual preprocessing is often required to achieve successful registrations with light-based microscopy images ^10^. Kim *et al*. ^5^ recently demonstrated that Otsu thresholding ^18^ —an algorithm to separate background from foreground pixels by maximizing inter-class variance—and maximizing pixel cross-correlation is a viable method for registering tissue slides imaged with IF and IMC, but it does not generalize in the absence of significant per-image manual tuning of preprocessing parameters. Compounding this issue is the lack of publicly available large datasets of same-slide multi-modal imaging on which to develop and evaluate registration workflows at scale.

To address this, we introduce Twocan, a workflow to search for a suitable cross-modality registration with minimal human intervention. Twocan leverages Bayesian optimization to select data preprocessing parameters to transform images that can be registered to maximize the number of registered nuclear pixels and cross-modality signal correlation **(Fig. 1A)**. This enables quick search and iteration through many parameter choices and register images for both cell line and tissue slides. We further introduce Twocan-954, the largest dataset of 954 paired IF and IMC images, including cell line and tissue slides under several different experimental conditions and microscopy settings as an evaluation resource for the broader community **(Fig. 1B)**, and use it to benchmark Twocan. Using Twocan-954, we demonstrate that no single preprocessing setting is sufficient to correctly register all images and that employing Twocan vastly improves the success of registering paired IF-IMC images across experimental conditions on both cell lines and tissues, outperforming both random baselines and existing methods.

**Figure 1.**
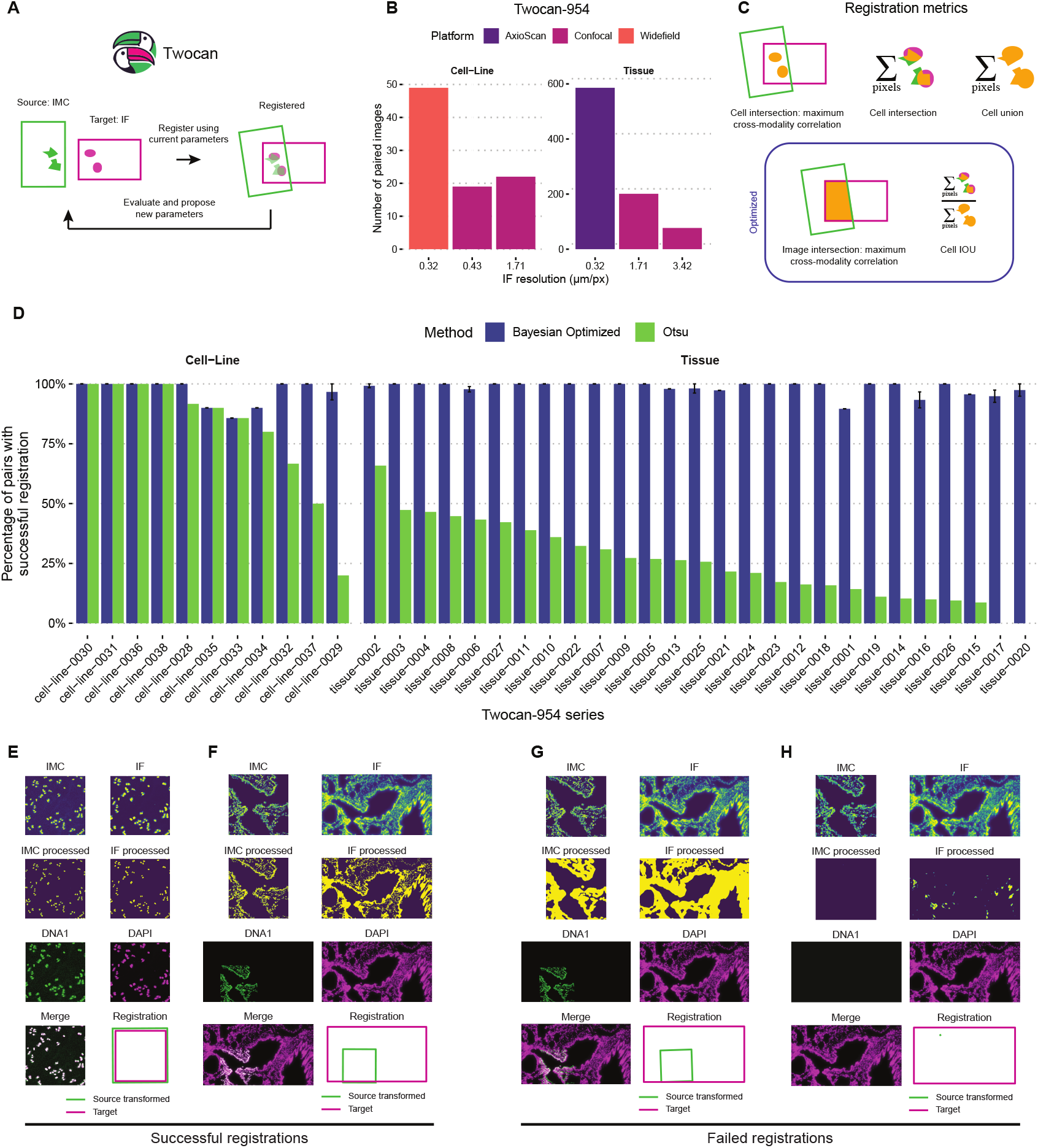
Methods overview. **(A)** For a given pair of images, Twocan iterates by proposing preprocessing parameters, and checking the resulting registration for the proposed settings. The parameter space is searched by optimizing registration quality metrics. **(B)** The Twocan-954 dataset contains 90 paired IF-IMC cell line images and 864 paired tissue images, imaged with a variety of microscopy platforms and resolutions. **(C)** Quality metrics used to evaluate registrations. **(D)** Bayesian optimized preprocessing parameters yield successful registrations more often than a Otsu-thresholding approach. **(E)** Example of processing and registration steps applied to a human osteosarcoma cell line U2OS. **(F)** Example of processing and registration steps applied to wildtype mouse lung tissue. **(G & H)**. Example of registration failures.

## RESULTS

### Development of a unique data resource of co-stained IF-IMC images

We first generated a comprehensive dataset of co-stained same-slide IF and IMC images **(Fig. 1B)**. This dataset—which we call Twocan-954—includes samples from three cell lines: U2OS (human osteosarcoma cell line), DLD1 (human colon carcinoma cell line), 293FT (transformed human embryonal kidney cell line), and lung tissue harvested from mouse (**Methods, Table S5**). The image pairs were collected under several experimental conditions to ensure broad applicability: untreated and wild-type controls, samples exposed to ionizing radiation, and cases with protein overexpression. To evaluate technical robustness of registration, we further varied multiple imaging parameters including magnification, size of field of view, and density of cells. All IF images were stained for nuclei with DAPI, and in all IMC panels included DNA targeting channels. Between three and 25 other channels were included within the IMC, which were held-out from registration and later used to evaluate performance. Slides were co-stained with fluorescent and metal-conjugated secondary antibodies targeting the same primary antibody. Subsequently, regions of interest imaged by IF microscopy were then captured with IMC laser ablation to attain images of identical regions on IF and IMC, allowing for rigid alignment between IF and IMC images. For each IF image, we selected up to five regions of interest for subsequent IMC analysis.

IF image dimensions were on average larger (mean 1050×1142 pixels) with resolution ranging from 0.32*µm*/px to 3.42*µm*/px, while IMC images were smaller (mean 338×347) with resolution 1*µm*/px. In cell line images, the IF had the same field of view as IMC, while in the tissue slides the IMC ranged from 1/100th to one third the field of view size of the IF. In total, we collected paired images from 90 cell line slides and 864 tissue slides, representing the largest dataset of paired IF-IMC images to date.

As there is no ground truth to determine whether two images are correctly registered, we next developed a set of metrics to measure the quality of a given candidate registration **(Fig. 1C)**. We quantify the total area of overlapping thresholded pixels which can be interpreted as a slide-specific absolute scale of nuclei overlap (`Cell intersection`) and the intersection over union of thresholded pixels gives a measure of the relative spatial overlap of nuclei in both modalities (`Cell IOU`). We also quantify the cross-modality correlation of nuclear channels across image modalities, which can be interpreted as measuring the signal agreement (`Cell-intersection correlation`) and the overall correlation within overlapping regions of both images (`Image-intersection correlation`). We created a multi-criteria filter for candidate registrations: (1) the proposed transformation matrix exists, and is invertible, (2) at least 100 pixels in the `Cell union` of processed pixels, (3) at least 0.5 `Image-intersection correlation`, and (4) at least 0.5 `Cell IOU`. A registration which met these criteria was deemed “successful” **(Fig. 1D,E,F)**. We found there was a wide range of parameter setting which returned a successful registration **(Fig. S1A)**.

Given these criteria, we investigated failure modes. **1**G illustrates a low-correlation registration, and 1H illustrates a representative example where the IMC image is transformed to be only a few pixels large, and compresses away all image signal. In this failure mode, individual metrics may be high; often `Image-intersection correlation` is equal to 1.0 **(Fig. S1B)** and this made up 32% of failure cases. Most (68%) failed trials were deemed failed because no transformation matrix was found **(Fig. S1C)**. This reflects a critical pain-point in the manual registration setting: most choices for preprocessing do not result in a pair of images that registration algorithms can automatically find landmarks to match on.

### Bayesian optimization for automated registration of immunofluorescence and multiplexed imaging

Current methods require intensive manual image pre-processing to enable registration of highly-multiplexed data. Therefore, to enable scalable and flexible registration we developed Twocan, a Bayesian optimization framework to automatically select the correct parameters to prepare a highly-multiplexed image, and IF image for registration. Bayesian optimization is a gradient-free strategy for function optimization, particularly useful when evaluations are costly – in this case, optimizing registration quality, when proposing a registration. By constructing a surrogate model to estimate both the optimal parameter settings and the associated uncertainty, this approach guides the selection of subsequent samples, balancing exploration and exploitation and is a common approach in processing biological data ^19^. Twocan first proposes a set of parameters to preprocess the images, including signal binarization threshold, level of gaussian blur, winsorization limits, whether to arcsinh transform the IMC data, and arcsinh cofactor (Table S1). After appropriately transforming the images, Twocan then searches for a rigid transformation which may include rotation, translocation, and scaling to maximize the signal correlation and agreement in cell location across modalities. Twocan then uses Optuna ^20^ to perform Bayesian optimization hyperparameter selection to find the per-image optimal parameter settings for registration **(Fig. 1A)**.

### Benchmarking using the Twocan-954 dataset

We applied our Bayesian approach across the entire Twocan-954 dataset, compared to a previously proposed Otsu-thresholding based method to register IMC and IF^5^. We found Twocan could register 96% of cell line slides compared to 82% using the Otsu approach. For tissue sections, this difference was even more pronounced, as Twocan could register 98% of the images Otsu could only register 28% **(Fig. 1D)**.

We found that the registrations returned by Twocan contain a wide range of preprocessing parameter settings. Surprisingly arcsinh-transformation, which is a common normalization to apply to single-cell segmented IMC data ^21^,^22^, is not always selected among the set of returned registrations (median 504/937 or 54% of successful registrations, three random seeds) **(Fig. S1A)**. We also found that IMC images were on average registered with a higher level of gaussian blur than IF, and that tissue slides were on average registered with lower IMC binarization threshold than cell-line slides **(Fig. S1A)**.

### Investigating search strategies for optimal registration

Having established that Bayesian optimization can reliably obtain successful registrations, we then sought to comprehensively characterize the optimal search and scoring strategies for Twocan. To do so, we compared six combinations of sampler and objective. The sampling methods we considered included (1) Gaussian Process (GP), which fits a GP to to the objective function and optimizes the acquisition function to suggest the next parameters, (2) Tree-structured Parzen Estimator (TPE), which fits two Gaussian mixture models to split the search space into parameters which lead to good performance, and those which lead to bad performance, and (3) Random, a baseline sampler which proposes new parameters uniformly at random. These samplers were used in combination with one of two objective functions for optimization: a multi-objective score incorporating Image Correlation and Cell IOU and a single-objective score which is the product of these two values.

To benchmark these search and scoring strategies, we selected a single successful IF/IMC registration for each strategy by ranking on the Balanced Score which balances `Cell intersection`, `Cell IOU`, and `Cell-intersection correlation`, normalized to the largest value for each metric (Methods, Figure S2A). This score was appropriate for selecting a registration to compare each strategy because it was not directly optimized by any of them and therefore allowed us to select a single best registration out of potentially many pareto-optimal solutions **(Fig. S2B)**. Once a best registration was returned for each Twocan sampler and objective combination, we evaluated the `Image-intersection correlation`of held-out channels which were not used for registration **(Fig. 2B,C)**.

**Figure 2.**
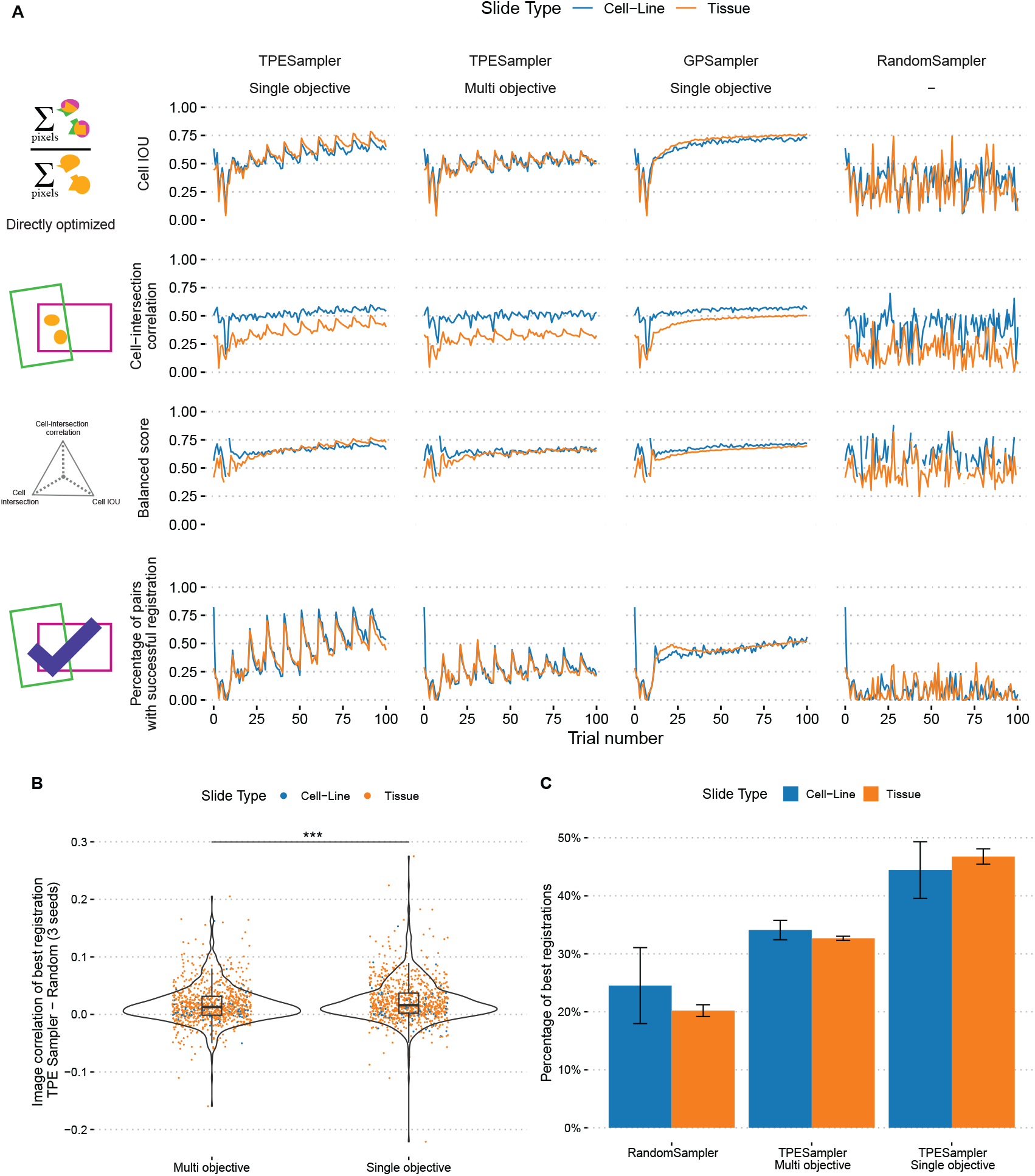
Comparison of three samplers and two objectives for an optimization budget of 100 trials across three seeds. **(A)** Registration metrics across 100 optimization trials (mean of 3 random seeds). **(B)** Difference of TPESampler vs Random for image cross-modality correlation on held-out channels. Boxes indicate median (center line), interquartile range (box), and whiskers extending to the most extreme points within 1.5x interquartile range. *** indicates t-test *P <* 0.001. **(C)** Percentage of pairs for which the best registration of each optimization strategy had highest cross-modality correlation on held-out channels. Bars indicate mean *±* standard error over three random seeds.

For both TPESampler and GPSampler, registration quality improved over the course of optimization, yielding higher `Cell IOU`, `Cell-intersection correlation`, Balanced score, and fewer registration failures. As was expected, this was not the case for the RandomSampler, which did not improve with increased optimization budget **(Fig. 2A)**. Both TPESampler and GPSampler found better registrations than the RandomSampler in terms of `Cell IOU`and `Image-intersection correlation`. Across the 954 image pairs, the best registrations were found with the TPESampler and single-objective strategy, which had the fewest failures overall **(Fig. S2C)** and achieved significantly higher `Cell IOU` (t-test, P*<*0.001) and `Cell-intersection correlation` (t-test, P*<*0.001) than Random sampling **(Fig. 2A)**.

Notably, the same transformation matrix may result in different values for those metrics which are defined over the pixels above a selected threshold depending on preprocessing. Consequently, lower thresholds may artificially inflate `Cell IOU`or `Cell-intersection correlation`, even when the underlying registration remains unchanged. This may incline the Bayesian optimizers to select for lower thresholds when it does not sacrifice registration success. We found that the best registration matrices for all successful registrations were similar to each other (within *±*15?pixels) regardless of choice of sampler or optimizer.

To compare different registrations even when they have highly similar transformation matrices, we next quantified the `Image-intersection correlation`of held-out channels not seen during registration. This independent metric was selected because it does not rely on any cell segmentation, thresholding, or other preprocessing parameter choices. The effect size of using TPESampler and single-objective approach compared to Random sampling was small (mean 0.023 improvement in Pearson correlation) but highly significant (t-test, P*<*0.001). This was also a significantly larger improvement over random compared to the TPESampler and multi-objective strategy (mean improvement in pearson correlation 0.017, **(Fig. 2B)**). When ranking by Image correlation of held-out channels and allowing for ties, TPESampler and single-objective most frequently found the best registration **(Fig. 2C)**.

For all Twocan setups, 9/954 pairs never achieved a successful registration, after the compute budget of 100 trials was exhausted **(Fig. S3)**. This is likely due to the IF image containing a region of over-exposed cells. Upon visual inspection it appears that some of these pairs were in fact well-registered, however they were deemed failures due to low correlation of the registration channels**(Fig. S3)**.

### Extending registration to additional modalities

We next investigated whether Twocan can register highly multiplex images with other modalities beyond protein-based IF. To do so, we retrieved two technical replicates of HeLa cell slides simultaneously imaged with RNA fluorescence in situ hybridization (FISH) and IMC from Schultz et al. 2018^23^. RNA-FISH allows visualization of individual mRNA transcripts, and these appear as bright spots of signal at sub-cellular resolution. Schultz et al. perform fluorescence microscopy followed by IMC on the same slides and registered in FIJI^24^ using the turboreg ^25^ plugin with manually selected landmarks, in order to compare the detection of PPIB in these two modalities and validate their simultaneous imaging technique. Following their analysis, we quantified spots of the housekeeping gene PPIB using Polaris ^26^ **(Fig. S2D)** and calculated the correlation of the number of RNA spots detected (mean 74 and 50 for replicates one and two respectively) to the total IMC ion counts per cell.

We ran Twocan to register these paired images and found spearman correlations of [0.90, 0.87] as compared to Schultz et al.’s manual registration which achieved spearman correlations of [0.88, 0.87]. Both the FISH binarization threshold, and the IMC binarization threshold parameters selected for best registrations were higher than the highest threshold selected in the IF and IMC from Twocan-954, while the remaining parameters fell within the same ranges as for the Twocan-954 dataset. This shows that Twocan can achieve at least human level performance, and even surpass it, without manual intervention and across a variety of multimodal technologies.

## DISCUSSION

Here we introduced Twocan-954, the largest and most comprehensive dataset of paired and registered IF and IMC images currently available **(Fig. 1)**, along with Twocan, a machine learning framework to register paired multi-modal spatial proteomics data with IF images without human supervision. We benchmarked several combinations of sampler and optimizers, and found that the best strategy is TPESampler and single-objective **(Fig. 2)**. Using this Twocan strategy, we successfully registered more than 98% of the images in our dataset by employing Bayesian optimization to select preprocessing parameters, significantly outperforming other approaches in consistency of both finding a suitable registration, and the registration quality.

A limitation of our analysis is that we restricted Twocan to rigid transforms (i.e., scaling, rotation and transformation only). This was appropriate for our dataset because we took a co-staining approach, which largely eliminates the possibility of distortions being introduced by rounds of serial staining or allowing slides to dessicate between imaging modalities ^23^. This is not the case in other common microscopy imaging settings such as for registering serial sections where a non-rigid transformation may be required ^10^. Additionally, we observed some examples of successful registrations which did not pass our successful registration filter **(Fig. S2)**, highlighting the challenge of establishing strict criteria to perfectly separate registration failures from successes.

We expect the Twocan-954 dataset will provide a comprehensive foundation for evaluating future registration methods for highly-multiplex and multimodal spatial datasets. This extensive community resource opens several exciting avenues of investigation. Firstly, the Twocan-954 dataset will support the development of emerging deep learning methods which leverage paired ‘ground truth’ from different technologies for tasks such as generating super-resolution microscopy ^27^, cell and tissue segmentation ^5^, or spot detection ^26^. Secondly, the adaptability of our Bayesian approach suggests that Twocan can be applied to a variety of imaging modalities beyond IF and IMC, for example other mass-spectrometry based approaches like MALDI, MSI, and potentially extended to spatial transcriptomics technologies like Xenium. Ultimately the ability to reliably integrate multi-modal datasets at scale, as demonstrated by Twocan, will empower more sophisticated downstream analyses to dissect complex cellular ecosystems and uncover novel biological insights previously obscured by modality-specific limitations ^8^. Together, the Twocan framework and the Twocan-954 dataset provide essential tools and resources that constitute a significant contribution to multi-modal spatial proteomics analysis.

## METHODS

### Cell culture and irradiation

U2OS (human osteosarcoma cell line), DLD1 (human colon carcinoma cell line) and 293FT (transformed human embryonal kidney cell line) cells were cultured in a sterile cell culture environment and were routinely tested for mycoplasma contamination. U2OS and 293FT cells were grown in Dulbecco’s modified Eagle’s medium (DMEM Wisent™, 319-015-CL), DLD1 cells were grown in RPMI-1640 (Wisent™, 350-015 CL). All cell culture medium was supplemented with 10% fetal bovine serum (FBS) and penicillin-streptomycin antibiotics, under standard cell culture conditions in CO2 incubators (37 °C, 5% CO2). U2OS cells carrying the FUCCI reporter, in which an exogenous fragment of the Geminin protein is tagged with the green fluorescent protein mAG and an exogenous fragment of the Cdt1 protein is tagged with the orange fluorescent protein mKO2, were generated as described ^28^. U2OS cells carrying a CDK2 sensor was generated as described ^29^, but with a modified construct in which an exogenous fragment of the DNA Helicase B protein is tagged with the red fluorescence protein mCherry and an exogenous fragment of the histone 2B protein is tagged with green fluorescent protein. Irradiation of cells was done in a Faxitron 43855D irradiator or a Precision X-Ray X-RAD® iR-160 irradiator.

### Immunostaining of cultured cells

Cells were grown on 3-well removable chamber slides (Ibidi 80381) and washed 2x with PBS before fixation with 4% PFA for 10 min at room temperature. Cells were washed 2x with TBS-0.1% Tween at room temperature, incubated with 100 nM Glycine for 10 min, then incubated with blocking buffer (10% horse serum + 3% bovine serum albumin in TBS-0.5% Tween) for 1 h. Cells were incubated with primary antibody (5 µg/ml each unless otherwise stated) in TBS-0.5% Tween at 4 °C overnight. (The primary antibodies used are described in the antibody panel above.) Cells were washed 3x 5 min with TBS-0.1% Tween, then incubated with fluorescent secondary antibodies only or a mix of fluorescent and metal-conjugated secondary antibodies (2 µg/ml each) in TBS-0.5% Tween for 1 h at room temperature in the dark. (The secondary antibodies used are described in the antibody panel above.) After washing 3x 5 min with TBS-0.1% Tween and once with TBS, cells were incubated for 5 min with 5 µg/ml 4’,6-diamidino-2-phenylindole dihydrochloride (DAPI; Thermo Fisher Scientific D1306). Cells were washed 2x 5 min with TBS, then mounted in TBS with 10% glycerol on a coverslip and imaged for fluorescence signal. The coverslip was then removed and cells were washed 2x 5 min with TBS, followed by a 5 min incubation with 500 nM Cell-ID Intercalator (Standard Biotools 201192B). Subsequently, cells were washed 2x 5 min with TBS, briefly dipped in water, dried, and subjected to mass cytometry laser ablation and acquisition.

### Immunostaining of mouse tissues

Tissue samples were formalin-fixed and paraffin-embedded at Mt Sinai Hospital, Toronto. Tissue sections were dewaxed by baking at 60 °C for 1 h followed by 3x 10 min washes in xylene, then rehydrated in a graded series of alcohol (ethanol:deionized water 100:0, 100:0, 96:4, 90:10, 80:20, 70:30; 5 min each). In a 95 °C water bath, heat-induced epitope retrieval was conducted in Tris-EDTA buffer at pH 9 for 30 min. The samples were blocked with 5% horse serum + 3% BSA in TBS-0.3% Triton for 1 h. Samples were incubated with primary antibody (5 µg/ml each unless otherwise stated) in TBS-0.5% Tween at 4 °C overnight. (The primary antibodies used are described in the antibody panel above.) Samples were washed 3x 5 min with TBS, then incubated with fluorescent secondary antibodies only or a mix of fluorescent and metal-conjugated secondary antibodies (2 µg/ml each) in TBS-0.5% Tween for 1 h at room temperature in the dark. (The secondary antibodies used are described in the antibody panel above.) After washing 3x 5 min with TBS-0.1% Tween and once with TBS, samples were incubated for 5 min with 2 µg/ml 4’,6-diamidino-2-phenylindole dihydrochloride (DAPI; Thermo Fisher Scientific D1306). Samples were washed for 15 min with TBS, then mounted in TBS with 10% glycerol on a coverslip and imaged for fluorescence signal. The coverslip was then removed and samples were washed 2x 5 min with TBS, followed by a 5 min incubation with 500 nM Cell-ID Intercalator (Standard Biotools 201192B). Subsequently, samples were washed for 15 min with TBS, briefly dipped in water, dried, and subjected to mass cytometry laser ablation and acquisition.

### Widefield and confocal microscopy

Images of fluorescent secondary antibodies, in addition to GFP and RFP fluorescence from U2OS-FUCCI reporter and U2OS CDK2 sensor cell lines, were acquired by slide scanner, widefield and confocal microscopy. Slide scan images were acquired on a Zeiss AxioScan1 slide scanner, equipped with a Orca Flash 4.0 V2 Axio Scan camera. A PlanApo 20X/0.8 objective was used for image acquisition. Widefield images were acquired on an Olympus BX61 upright fluorescence microscope, equipped with a Hamamatsu DP71 camera. A UPlanSApo 20X/0.75 air objective was used for image acquisition. Confocal images were acquired with a Nikon A1 confocal laser scanning microscope. 10x and 20x air objectives were used for image acquisition. The sequential scanning mode was applied, and the number of overexposed pixels was kept at a minimum. Z sections were recorded with optimal distances based on Nyquist criterion. Imaging mass cytometry.

Images were acquired using a Hyperion Imaging System (Fluidigm). Fluorescence images acquired prior were compared to brightfield scans of the cultured cells or tissues on the Hyperion Imaging System to identify regions of alignment to be acquired. Rectangular areas from stained cultured cells and tissue samples were laser ablated in a rastered pattern at 200 Hz, and raw data preprocessing was completed using commercial acquisition software (Fluidigm).

### Data preparation

Fluorescence and IMC images were converted to .tiff format. For fluorescence images acquired as Z sections, maximum intensity projections were calculated using Fiji ^24^. IMC images acquired in .mcd format were converted to .tiff format using the readimc package ^30^. IF images for each protein were converted to greyscale, and concatenated into a single multi-channel TIFF. For each image series, channels were selected as “registration channel” and “evaluation channel” as listed in Table S2. The preferred choice for registration channels is DNA or nuclear-localizing proteins.

### IF preprocessing

Preprocessing parameters are proposed with Optuna V3.6.1, using one of: GPSampler, TPESampler, Random sampler. Following Muhlich et al. ^16^ IF images are rescaled by the square root of their resolution. Then, registration channels are summed across the channel axis and scaled by the maximum value in the image. A gaussian blur filter is then applied with the Optuna-proposed variance. Then the image is binarized according to the proposed threshold.

### IMC preprocessing

Preprocessing parameters are proposed with Optuna, using one of: GPSampler, TPESampler, Random sampler. Whether to arcsinh transform the data is proposed, and if so then IMC data is transformed according to the proposed arcsinh cofactor. Then, registration channels are summed across the channel axis, and the signal is winsorized according to the proposed winsorization limits. The image is then scaled by its maximum value and a gaussian blur filter is applied with the proposed variance. Then the image is binarized according to the proposed threshold.

### Registration metrics

We evaluated registration quality using several complementary metrics that assess both spatial alignment and signal correlation.

#### Spatial overlap metrics

The spatial overlap between IF and IMC images was assessed using binary masks of cell-containing regions. We computed:

- `Cell intersection`: The number of pixels above the set thresholds in both modalities
- `Cell union`: The number of pixels above the set thresholds in either modality
- `Cell IOU`: The ratio of intersection to union, providing a normalized measure of overlap between 0 and 1

#### Correlation metrics

We computed correlations between registered images at two levels:

- `Cell-intersection correlation`: The maximum Pearson correlation coefficient between any pair of channels across modalities, calculated only over pixels above the set thresholds in both modalities.
- `Image-intersection correlation`: The maximum correlation coefficient calculated over all pixels in the overlapping field of view.

For both correlation metrics, we computed three variants:

- Stack correlation: Considering all available channels
- Registration correlation: Considering only the channels used for registration
- Evaluation correlation: Considering only held-out channels not used in registration

All correlations were computed after registration and rescaling of the IF image to match IMC resolution. To ensure robust comparison, correlations were calculated only over the intersection of the transformed images, avoiding edge artifacts and regions without data in either modality.

### Criteria for a successful registration

The following criteria were used to ensure a minimum quality on registrations:

- The proposed transformation matrix exists, and is non-singular
- At least 100 pixels in the `Cell union`of processed pixels
- At least 0.5 `Image-intersection correlation`
- At least 0.5 `Cell IOU`

### Image registration

Twocan by default is designed to register IMC images onto IF, and is flexibly adaptable to other modalities, and we leave open the option for custom preprocessing functions. For both the IF and IMC, proposed preprocessing parameters are used to create a “registration image”. These registration images are no longer multiplexed, and are registered using ORB keypoint detection and matching. We use OpenCV V4.10.0 for this, and return an estimated affine transformation matrix, M. For each registration trial, if M exists and is non-singular, then we apply it to the registration channels of the original image and score the quality of the registration with the previously described metrics. For the first optimization trial, we enqueue parameters equivalent to Kim *et al*. ^5^: Otsu thresholding for both IF and IMC, 0 gaussian blur, no arcsinh transformation and winsorization limits of 1%. After this we use the Optuna defaults for Bayesian samplers to have a burn-in of 10 random trials.

### Optimization objectives

We implemented two optimization strategies to identify optimal preprocessing parameters: a single-objective and a multi-objective approach. If no registration matrix is found, the objective values are set to 0. Both strategies combine Image correlation and Cell IOU metrics to evaluate registration quality. The theoretical minimums for these values are -1 and 0 respectively. The single-objective function multiplies the maximum correlation coefficient across registration channels with the IOU of the binary cell masks. This composite score rewards solutions that achieve both high spatial correlation and substantial cell overlap. The multiplication ensures that both criteria must be satisfied for a high score, as either poor correlation or minimal overlap will result in a low overall value.

The multi-objective function treats `Image-intersection correlation`and `Cell IOU`as separate optimization targets, allowing exploration of the trade-off between these metrics. This approach enables identification of Pareto-optimal solutions where neither metric can be improved without degrading the other. Optuna-based hyperparameter selection demonstrated improved stability when using bounded objective functions (e.g., [0,1] for intersection-over-union or [-1,1] for correlation coefficients). Optimization of unbounded metrics such as pixel counts proved challenging, possibly due to numerical instabilities in posterior estimation when handling failed registrations. While these issues could be addressed by setting failed registration scores to historical minima, we ultimately selected bounded metrics (Pearson correlation and intersection-over-union) which did not require such strategies.

### Choosing a single best registration

While we optimize on cross-modality Cell correlation and Cell IOU during registration, these metrics may be inflated when a high preprocessing threshold is selected, or in other words when the absolute number of pixels in the cell intersection is small. To address this, we developed a geometric scoring approach which additionally incorporates maximizing the Cell intersection. We included Cell Intersection in the Balanced Score because a higher value corresponds to matching on more of the image signal across both modalities. Additionally, Cell Intersection is measured in number of pixels, and depends both on the dimensions of the images and the relative fields of view. This makes it less natural to include in an objective function and is particularly valuable since it allows us to evaluate registration quality without relying on cell segmentation, which can have highly differing performance across imaging technologies and tissue density.

The Balanced Score is the area formed by these three normalized metrics as vertices in a unit simplex, where poor performance in any single metric reduces the overall score. Each metric was first normalized within each image pair to account for variation in absolute values across different samples:

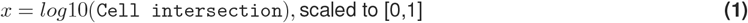

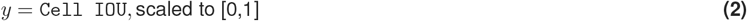

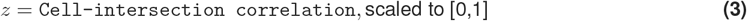

The balanced score is then calculated as ?*xy* ±?*yz* ±?*xz*?*/*3?The maximum possible value is 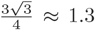 which is the area of the equilateral triangle with vertices 1 unit away from the origin **(Fig. S2A)**. The balanced score is a relative score, and is normalized within all successful registration trials proposed for each sampler and objective for each pair. This is appropriate for ranking proposed registrations for a pair of images, but does not give an absolute measure of registration quality.

## RESOURCE AVAILABILITY

### Lead contact

Requests for further information and resources should be directed to and will be fulfilled by the lead contact, Kieran R. Campbell.

### Materials availability

#### Data and code availability

Data and code required to reproduce this study can be retrieved from:

- Zenodo https://zenodo.org/records/15115811
- Github https://github.com/camlab-bioml/twocan

## ACKNOWLEDGMENTS

This work was supported by funding from CIHR project grant PJT175270 (KC), NSERC Discovery grant RGPIN-2020-04083 (KC) and DGECR-2021-00366 (HWJ), a CFI/JELF awards (KC, HWJ). This research was undertaken, in part, thanks to funding from the Canada Research Chairs Program. This work is supported by funding from the Data Sciences Institute, University of Toronto.

## AUTHOR CONTRIBUTIONS

Conceptualization, C.F.H., K.R.C., C.Y.L. and H.W.J.; Methodology, C.F.H. and K.R.C.; Software, C.F.H.; Formal Analysis, C.F.H.; Investigation, C.Y.L.; Resources, C.Y.L., D.C, C.L., R.B., H.W.J.; Data Curation, C.F.H.; Writing - Original Draft, C.F.H.; Writing - Review & Editing, C.F.H., K.R.C., C.Y.L and H.W.J.; Visualization, C.F.H.; Supervision, H.W.J. and K.R.C.; Funding Acquisition, C.F.H., H.W.J. and K.R.C.;

## DECLARATION OF GENERATIVE AI AND AI-ASSISTED TECHNOLOGIES

During the preparation of this work, the author(s) used claude.ai in order to edit text, and summarize code functionality. After using this tool or service, the author(s) reviewed and edited the content as needed and take(s) full responsibility for the content of the publication.

## SUPPLEMENTAL INFORMATION INDEX

Figures S1-S3 and their legends in a PDF

Tables S1-S5 in an excel file

