## supplemental figures for "Automated registration of spatial expression data scales multimodal integration to large cohorts"

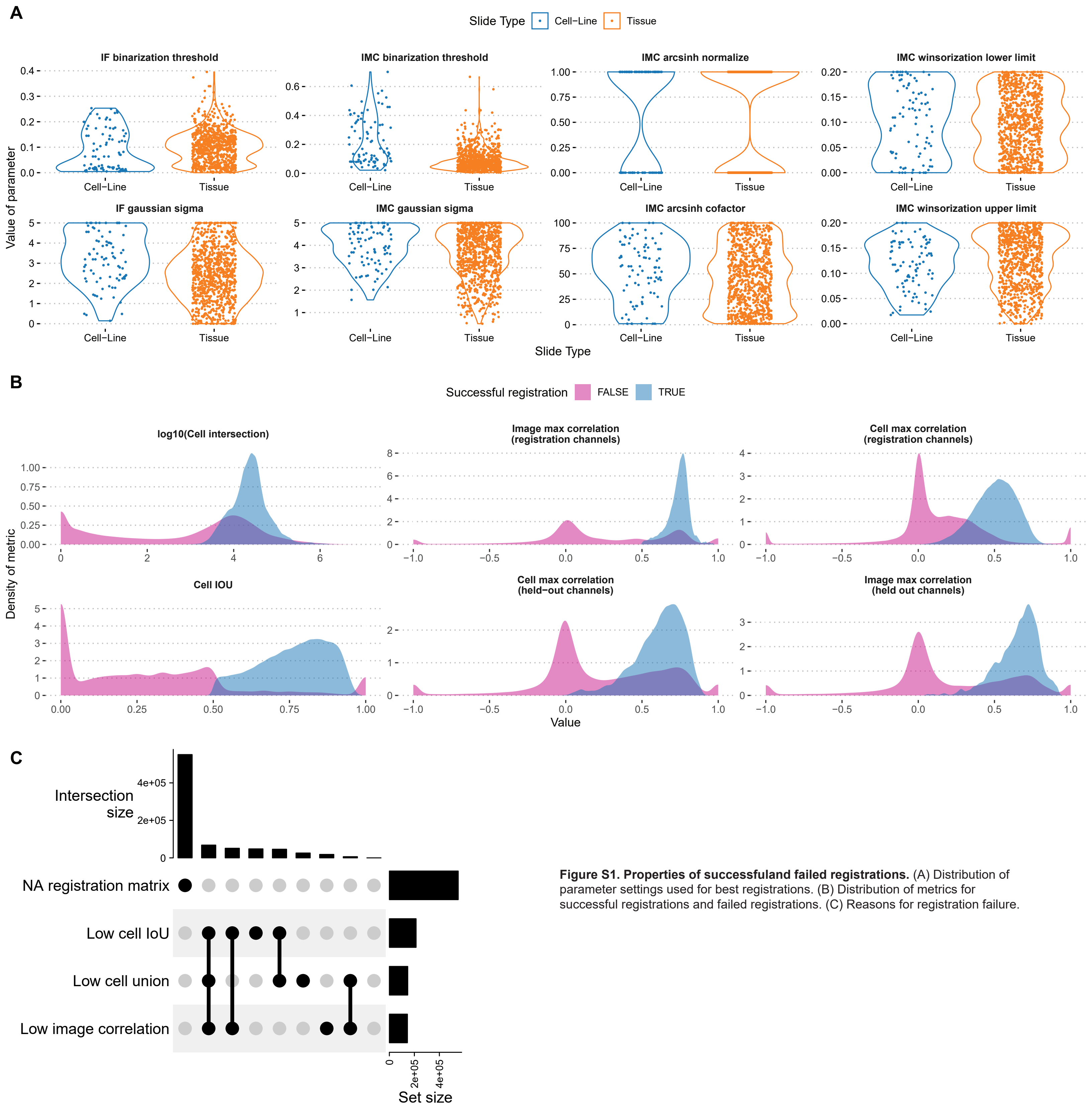


**Figure S1. Properties of successfuland failed registrations.** (A) Distribution of parameter settings used for best registrations. (B) Distribution of metrics for successful registrations and failed registrations. (C) Reasons for registration failure.

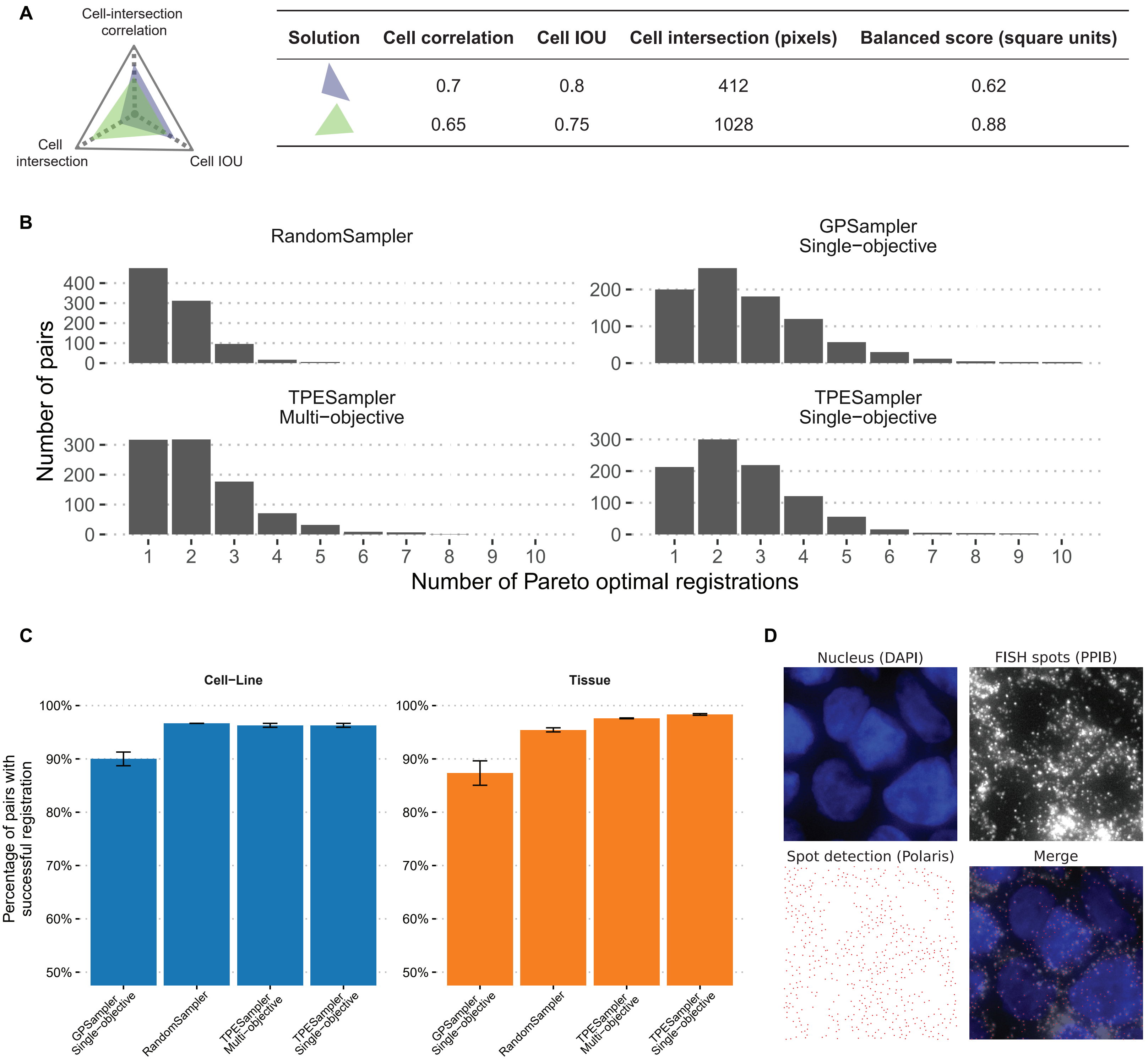


**Figure S2. Comparison of Twocan strategies combining different samplers and objectives.** (A) Geometric interpretation of the Balanced Score. (B) Distribution of the number of registrations found by four Twocan strategies which are pareto optimal with respect to the Cell IOU and Cell-intersection correlation. (C) Frequency of registration success for each Twocan strategy. (D) Example of FISH spots detected by Polaris.

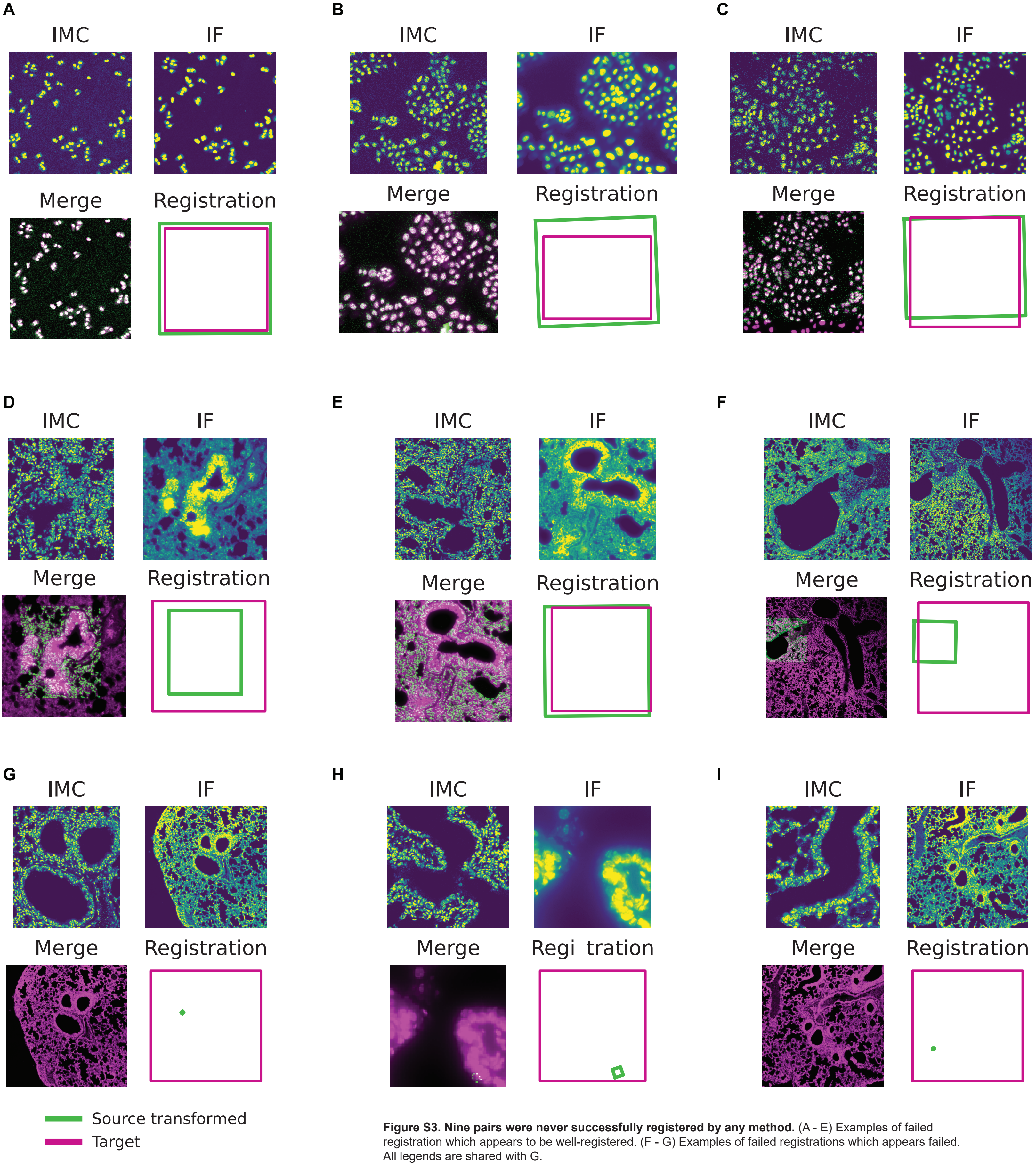


**Figure S3. Nine pairs were never successfully registered by any method.** (A - E) Examples of failed registration which appears to be well-registered. (F - G) Examples of failed registrations which appears failed. All legends are shared with G.
